# Expanding *Ventilago* to incorporate *Smythea*: new combinations in Ventilagineae (Rhamnaceae) based on novel sequence data and phylogenetic analysis

**DOI:** 10.1101/2025.03.26.645417

**Authors:** Henry Miller, Daniel Cahen, Felix Forest, Nattanon Meeprom, Pirada Sumanon, Timothy M. A. Utteridge

## Abstract

Ventilagineae (Rhamnaceae) is a tribe of palaeotropical climbing plants, comprising two genera: *Smythea* and *Ventilago*. Previous studies have established the monophyly of the tribe, but the relationships within it remain unclear. This research includes six species of *Smythea* and seven species of *Ventilago* in a molecular phylogeny using the Angiosperms353 probe set. The results show that *Ventilago* is paraphyletic, with *Smythea* nested within it. To resolve this issue, we propose to merge *Smythea* into *Ventilago*. To achieve this, we make seven new combinations in *Ventilago* as well as resurrecting five names previously published in *Ventilago* but subsequently transferred to *Smythea*. In addition, we typify *Ventilago*.

## Introduction

The buckthorn family Rhamnaceae contains c. 65 genera and over 1,200 species (WCVP 2024). Growth habits are diverse, ranging from herb to shrubs, climbers and trees (Richardson *et al*. 2000b). The tribe Ventilagineae within the Rhamnaceae includes only two genera, *Smythea* Seem. and *Ventilago* Gaertn. These genera contain 12 and 40 species, respectively, widely distributed in the Old-World tropics, from Africa to the Pacific Islands (Cahen 2022). *Ventilago* was established by Joseph Gaertner in 1788 (Gaertner 1788), based on specimens from the collection of Joseph Banks of *Ventilago madraspatana* Gaertn. and on *Funis viminalis* described by Rumphius (1747). The generic name probably refers to the wind-dispersed winged fruits, from the Latin *ventus* (wind) (Cahen & Utteridge 2017a). *Smythea* was published by Berthold Carl Seemann in 1862, based on specimens from Fiji. The generic name honours William Smythe (1816–1887), a British naval officer who undertook a mission to Fiji. Both genera, except for one species, *V. viminalis,* are climbers at maturity. Tropical climbing plants are some of the most under-studied species in the plant kingdom (Putz & Mooney 1991).

Modern classifications of Rhamnaceae reviewed by Cahen & Utteridge (2017a), base generic delimitations on seed chamber shape, with other characters sometimes being indicative **(see Figures 1a and b**) *Ventilago* was defined as having globose seed chambers, while *Smythea* has seed chambers which taper gradually into the wing, a character first suggested by Weberbauer (1895). *Smythea* and *Ventilago* were placed as the sole members within the tribe Ventilagineae Hook.f. (Hooker 1862). Members of the tribe are easily distinguished from other Rhamnaceae by the presence of a pronounced apical fruit appendage, in the form of a wing or inflated chamber.

**Figure 1a:**
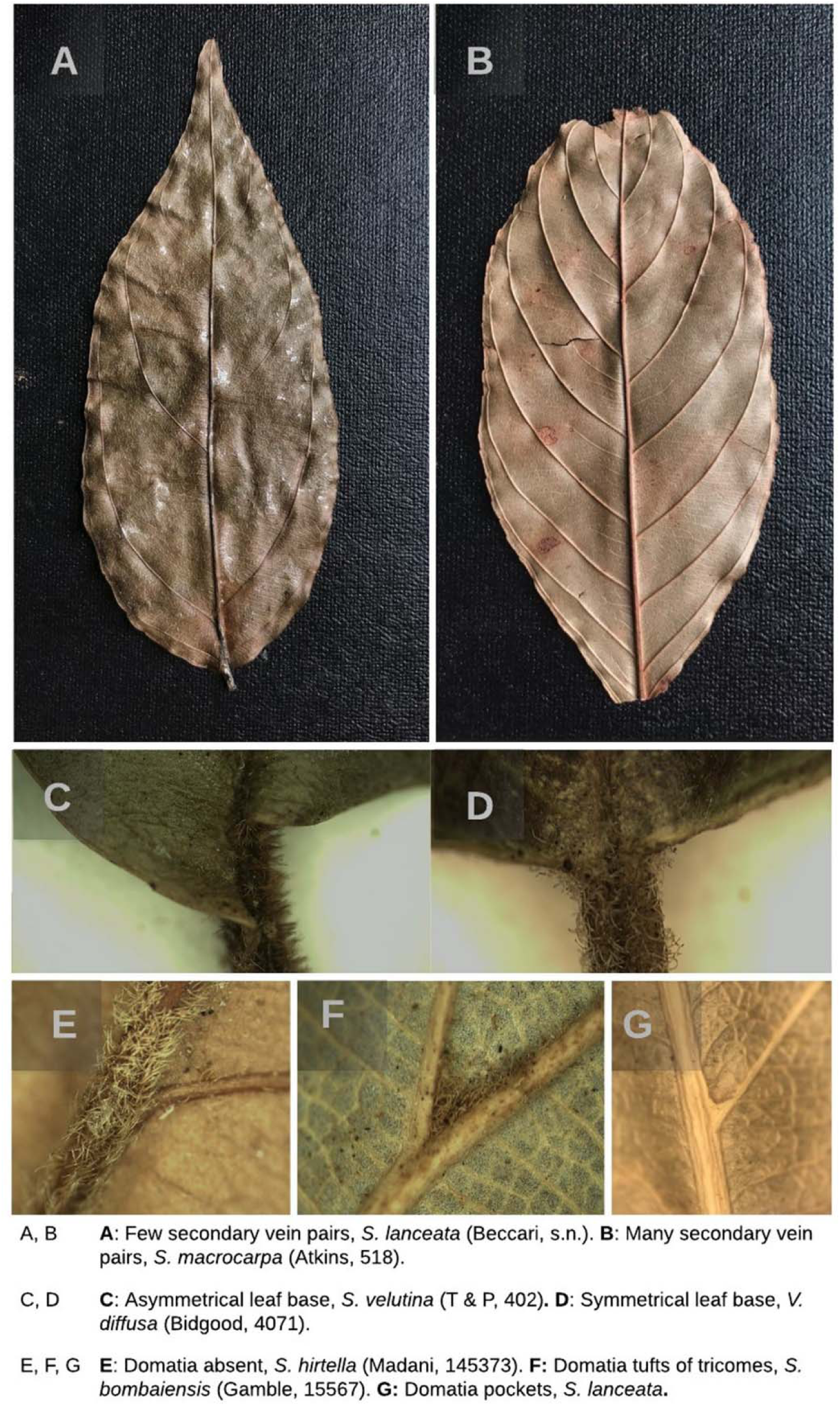
Variation in vegetative morphology used for generic delimitation in Ventilagi-neae.

**Figure 1b:**
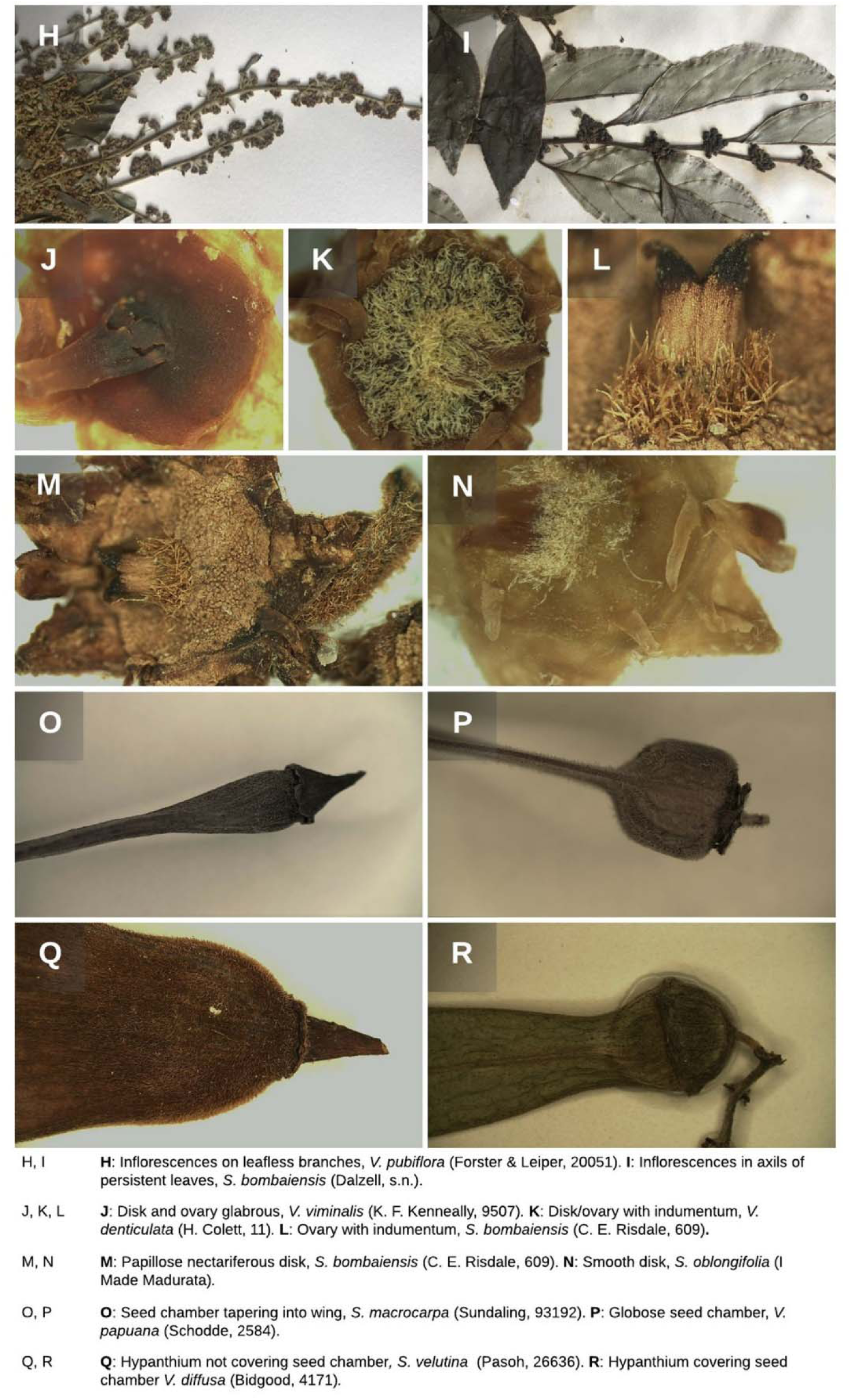
Variation in reproductive structures in Ventilagineae used for generic and specific delimitation.

The first family-wide molecular analysis of Rhamnaceae by Richardson *et al*. (2000a) revealed the existence of three well-supported clades: the ziziphoids, rhamnoids and ampeloziziphoids. Subsequent research has supported these results. Notably, Hauenschild *et al*. (2016) generated a phylogeny based on *trnL-trnF* and ITS markers, incorporating over 400 taxa and 57 genera, which further corroborated the monophyly of these three clades Tian *et al*. (2024) conducted an even more extensive sampling of Rhamnaceae, including 574 species (representing 52.2% of species and 92.0% of genera), and successfully sequenced 15 *Ventilago* and two *Smythea* species. Their maximum likelihood tree, published as this manuscript was in preparation, placed *S. lanceata* (Tul.) Summerh. and *S. oblongifolia* (Blume) Cahen & Utteridge as sister taxa nested within *Ventilago*. These phylogenetic analyses consistently place Ventilagineae within the rhamnoids as a sister group to the rest of the clade, which consists of *Maesopsis* Engl., *Fenghwaia* G.T.Wang & R.J.Wang, and the tribe Rhamneae Hook. f. (Richardson *et al*. 2000a, 2000b; Wang *et al*. 2021). It is important to note that Richardson *et al*. (2000a) did not include *Smythea* in their molecular analysis of Rhamnaceae, as they could not obtain suitable DNA samples from this genus. They tentatively placed *Smythea* within Ventilagineae based on its morphological similarity to *Ventilago*.

This research aims to improve our understanding of the relationships between these genera using molecular methods. We include half of the described species of *Smythea* in a molecular phylogenetic study with several *Ventilago* species using a targeted enrichment approach and the Angiosperms353 probe set (Johnson *et al*. 2018). Our objectives are to evaluate the generic delimitations of *Smythea* and *Ventilago*, identify potential morphological diagnostic characters for the clades obtained, infer possible evolutionary events, and discuss the implications for conservation. Ultimately, we aim to resolve some of the taxonomic and phylogenetic uncertainties in Ventilagineae and establish a robust framework for future studies of this group.

## Methods: Molecular phylogenetics

### Sampling and DNA extractions

Leaf samples were taken from herbarium sheets at Kew (K). DNA was extracted using the CTAB extraction protocol (modified from Doyle and Doyle 1987). DNA was precipitated in isopropanol at −20°C for two weeks prior to resuspension and PCR.

### High-throughput sequencing

Nuclear genes enrichment was carried out using the Angiosperms353 probe set (Johnson *et al*. 2018). DNA was fragmented to approximately 350 bp using ME220 Focused-ultrasonicator (Covaris). The shearing step was omitted for some herbarium samples as their DNA was already fragmented to the desired size. The sheared DNA was developed into the DNA libraries suitable for Illumina sequencing using NEBNext Ultra II DNA Library Prep Kit for Illumina (New England BioLabs). The DNA libraries were indexed using NEBNext Multiplex Oligos for Illumina (Dual Index Primers Set) (New England BioLabs) and amplified using 8 PCR cycles. Next, the DNA libraries were quantified using Quantus Fluorometer (Promega) for yield and 4200 TapeStation System (Agilent) for library fragment size distribution and average library size. The indexed DNA libraries were equimolarly pooled and hybridised with the Angisperms353 probe set. The hybridised libraries were captured using a magnetic separator then amplified using the universal PCR primers for 14 PCR cycles. DNA libraries were sequenced using Illumina HiSeq X for 150 PE to ∼1Gbp per sample at Macrogen (South Korea).

### Phylogenetic reconstruction

For the reconstruction of phylogenetic relationships, we followed the framework provided by Sumanon *et al*. (2023). Raw sequence data was trimmed to remove adapters and low-quality reads using Trimmomatic v.0.39 (Bolger *et al*. 2014) with the MAXINFO algorithm with strictness 0.5 and reads shorter than 36 bp were eliminated. Read quality was assessed before and after trimming using FastQC v.0.11.9 (Andrews 2010). The HybPiper pipeline (Johnson *et al*. 2016) was applied, first using BWA v.0.7.17 (r1188) to map cleaned reads to the Angiosperms353 target sequences using the mega353 target file (McLay *et al*. 2021). Subsequently, on-target reads were assembled to each locus with SPAdes v.3.13.0 (Bankevich *et al*. 2012). Potential paralogs were detected by the script implemented in HybPiper and excluded from downstream analyses. Supercontigs which comprised exons targeted by Angiosperms353 and their adjacent splash zones were retrieved and used for downstream analyses.

Supercontigs of each locus were aligned using MAFFT v.7.475 (Katoh & Standley 2013) with local pairwise alignments, 1,000 iterative refinements and reverse complementation of sequences if necessary. Fragmentary sites of the alignments were removed using optrimAl script (Shee *et al*. 2020). Then the trimmed alignments were inspected and further cleaned manually in an alignment editor program. Preliminary gene trees were then generated using IQ-TREE v.2.0.3 (Minh *et al*. 2020) to detect long branches using TreeShrink v.1.3.7 (Mai & Mirarab, 2018). These long branches indicated possible sequence errors, thus were removed from the alignments.

Gene tree estimation was performed in IQ-TREE v.2.0.3 (Minh *et al*. 2020) using the cleaned/filtered alignments from the previous step. We set the parameter to do 1,000 ultrafast bootstrap replicates and used joint model testing in ModelFinder (Kalyaanamoorthy *et al*. 2017). Each gene tree was rooted with *Maesopsis eminii* Engl. using phyx v.1.3 (Brown *et al*. 2017) and internal branches which had bootstrap support below 30% were collapsed using Newick utilities v.1.6 (Junier & Zdobnov 2010). *M. eminii* was chosen as the outgroup because it is in the sister tribe to *Ventilago*, the Maesopsideae (Richardson *et al*. 2000a) and sequences were available online. Then a species tree was inferred from all individual gene trees using the coalescent-based method ASTRAL v.5.7.5 (Zhang *et al*. 2018).

Raw sequence data are deposited in Sequence Read Archive (SRA) available on NCBI servers under BioProject number PRJNA1100519 (http://www.ncbi.nlm.nih.gov/bioproject/PRJNA1100519). All scripts related to phylogenetic analyses follow Sumanon *et al*. 2023 and are available on GitHub at https://github.com/pebgroup/Maesa_sptree. The alignments, gene trees, and species trees are deposited on Zenodo at 10.5281/zenodo.10976870.

## Results

### Molecular phylogenetic tree

#### Sequencing results

– we retrieved between 2,212,034 and 21,531,577 trimmed reads per sample. A median of 64.6% were on target across all samples (see Table 1). After assembling the reads to the target reference, the median of loci retrieved per sample was 349. However, after alignment cleaning and filtering out paralogs and suspicious sequences, 343 alignments remained for downstream analyses. The supercontig alignments used in tree building had a total length of 734,653 bp covering 17 samples with 16.7% missing data. Of the total length, 17.6% and 6.8% were variable and parsimony informative, respectively. The statistics for 343 individual gene alignments are summarised and reported in Table 2.

**Table 1:** Summary statistics from targeted sequencing for 16 samples of *Ventilago* and *Smythea,* including *Maesopsis eminii* as an outgroup.

|  |  | Reads total | Reads on target |  | Number of recovered exons longer than X% of the target sequence length |  |  |  |
| --- | --- | --- | --- | --- | --- | --- | --- | --- |
|  |  | # | # | % | X = 0 | X = 25 | X = 50 | X = 75 |
| *Ventilago*<br>and *Smythea* | Minimum | 2,212,034 | 1,656,035 | 44.3 | 345 | 337 | 300 | 204 |
|  | Q1 | 9,017,765 | 5,393,937.75 | 59.8 | 347 | 342 | 313 | 230.5 |
|  | Median | 11,016,452.5 | 7,286,305 | 64.6 | 349 | 346 | 331 | 278.5 |
|  | Q3 | 14,396,808.25 | 8,586,502.25 | 68.25 | 350.25 | 348 | 336.25 | 289 |
|  | Maximum | 21,531,577 | 14,312,007 | 75.8 | 352 | 350 | 342 | 302 |
| Outgroup |  | 3,347,812 | 1,120,764 | 33.5 | 124 | 66 | 31 | 16 |

**Table 2:** Summary statistics for 343 alignments used in gene tree reconstruction and species tree inference.

|  |  | Median | Mean | SD | Min | Max |
| --- | --- | --- | --- | --- | --- | --- |
| Number of taxa |  | 17 | 16.9 | 0.6 | 10 | 17 |
| %Occupancy |  | 100 | 99.2 | 0.04 | 58.8 | 100 |
| Supercontigs | Alignment length | 1909 | 2141.8 | 1110.7 | 487 | 7449 |
|  | %missing data | 15.3 | 16.5 | 6.2 | 5.8 | 37.5 |
|  | Number of variable sites | 339 | 386.2 | 205.8 | 73 | 1394 |
|  | %variable sites | 17.6 | 18.1 | 3.3 | 10.2 | 32.2 |
|  | Parsimony-informative sites | 138 | 149.5 | 85.9 | 26 | 617 |
|  | %parsimony-informative sites | 6.8 | 6.9 | 1.7 | 3.5 | 16.7 |
| Exons | Alignment length | 651 | 738.4 | 434.2 | 91 | 3606 |
|  | %missing data | 5.6 | 7.9 | 7.5 | 0 | 43.2 |
|  | Number of variable sites | 100 | 112.9 | 71.4 | 9 | 502 |
|  | %variable sites | 14.5 | 15.2 | 3.6 | 4.4 | 31.2 |
|  | Parsimony-informative sites | 28 | 32.8 | 22.8 | 2 | 172 |
|  | %parsimony-informative sites | 4.2 | 4.3 | 1.5 | 0.8 | 13.6 |
| Flanking regions | Alignment length | 1244 | 1403.4 | 788.8 | 230 | 4328 |
|  | %missing data | 20.4 | 21.1 | 6.1 | 9 | 40.7 |
|  | Number of variable sites | 242 | 273.3 | 159.8 | 40 | 1086 |
|  | %variable sites | 18.8 | 19.6 | 4.1 | 10.2 | 39.2 |
|  | Parsimony-informative sites | 105 | 116.7 | 72 | 6 | 445 |
|  | %parsimony-informative sites | 8 | 8.3 | 2.2 | 0.02 | 0.2 |

#### Tree topology and branch support

**–** A well-resolved and well-supported molecular phylogenetic tree of Ventilagineae was obtained, based on target enrichment sequencing using the Angiosperms353 probe set and from the coalescent-based species tree inference (Figure 2). The species trees, inferred from 343 gene trees, included 82% of the quartet tree induced from the gene trees. Of the nodes in the species tree, 79% were well supported (local posterior probability, LPP ≥ 0.95), and the average LPP for the nodal support was 0.97.

**Figure 2:**
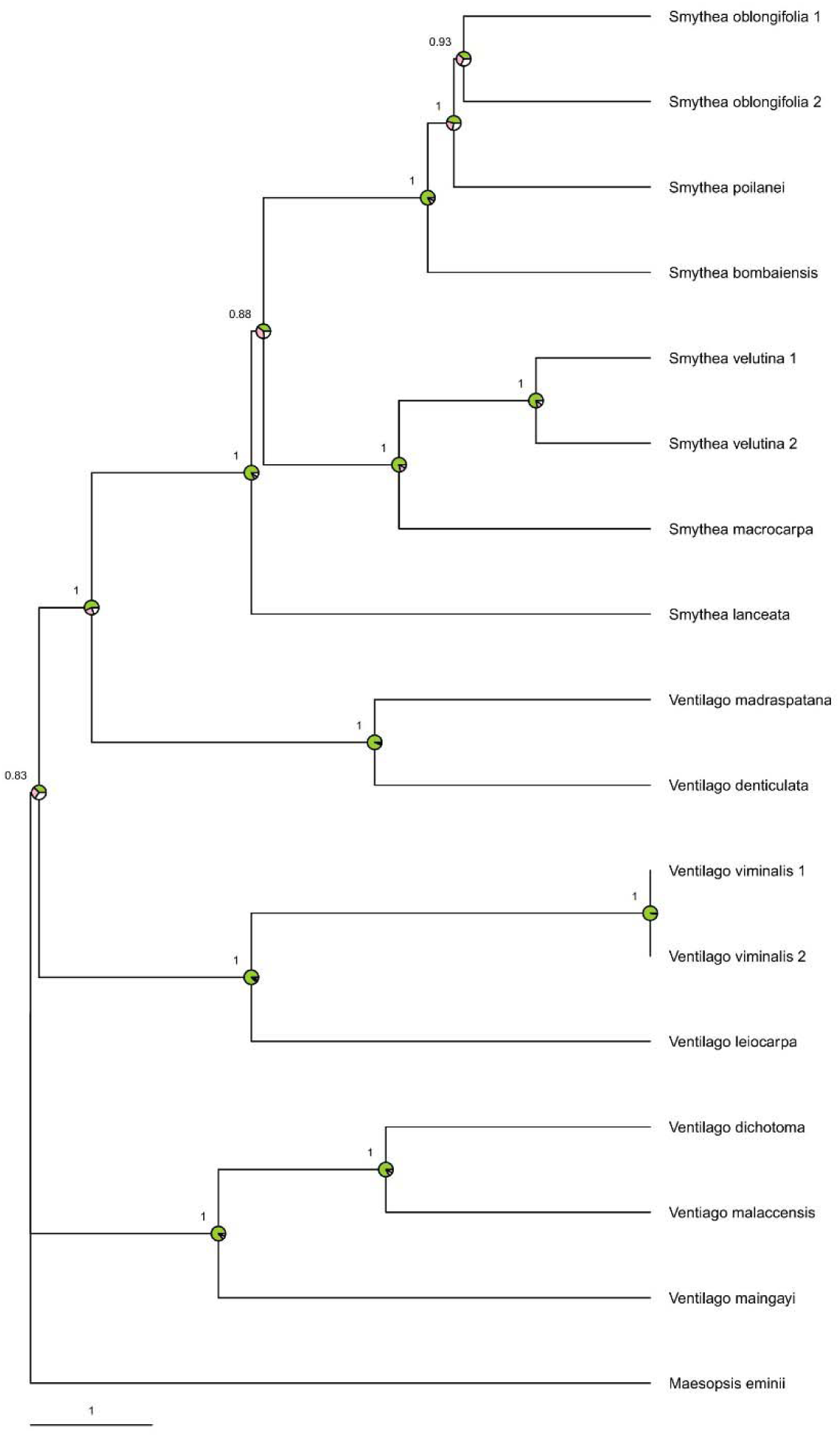
Phylogenetic tree of Ventilagineae using the ASTRAL-III coalescent algorithm of 353 nuclear gene trees. The support values shown are the Local Posterior Probability

The phylogeny shows with high support that *Ventilago* is paraphyletic and *Smythea* is monophyletic, nested within *Ventilago*. Six clades within Ventilagineae were identified, each containing only *Smythea* or *Ventilago* species: Clade A (*S. oblongifolia*, *S. poilanei* Cahen & Utteridge, and *S. bombaiensis* (Dalzell) S.P.Banerjee & P.K.Mukh.); Clade B (*S. velutina* (Ridl.) S.P.Banerjee & P.K.Mukh. and *S. macrocarpa* Hemsl.); Clade C (*S. lanceata*); Clade D (*Ventilago, V. denticulata* Willd. and *V. madraspatana* Gaertn.); Clade E (*V. viminalis* Hook. and *V. leiocarpa* Benth.); and Clade F (*V. malaccensis* Ridl., *V. dichotoma* (Blanco) Merr., and *V. maingayi* M.A.Lawson).

Within the well-supported Clade A, *S. oblongifolia* and *S. poilanei* are sister taxa, sharing several morphological characters: seed chambers that taper gradually into the wing, a calyx covering only a small portion of the seed chamber, and domatia composed of tufts of trichomes at the midrib/primary vein intersections. The most obvious differences are the much denser indumentum on the fruit, and the flower fascicles in leafless racemes or panicles in *S. oblongifolia* (vs in axils of persistent leaves in *S. poilanei*). *Smythea oblongifolia* and *S. poilanei* are sister to *S. bombaiensis*. These three species share the fruit and domatia characters namely tapering seed chambers and domatia composed of tufts of trichomes, but differ in *S. poilanei* having a less hairy fruit than the other two species, and *S. oblongifolia* having flower fascicles in leafless racemes or panicles (vs in axils of persistent leaves in *S. bombaiensis* and *S. poilanei*). *Smythea oblongifolia* also differs in the narrower angle of divergence between the secondary and primary veins, as measured on the apical side of the point of branching, 20 – 35° (vs 35 – 55°).

Clade B (*S. velutina* and S*. macrocarpa*) is sister to Clade A. The species of Clade B share fruit and domatia characters with Clade A. This includes seed chambers that gradually taper into the wing and domatia composed of tuft of trichomes at the midrib and primary vein intersections. In addition, *S. bombaiensis and S. oblongifolia* (Clade A) and *S. macrocarpa* and *S. velutina* (Clade B), have flowers with the disk surface sometimes covered by distinct cylindrical projections, a character absent from all other Ventilagineae examined. No flowering specimen of *S. poilanei* is available to determine whether this character could be diagnostic of both Clades A and B. Shorter papillae were also observed on the disks of some flowers of *S. beccarii*, but this species was not included in the current study. *Smythea velutina* and *S. macrocarpa* differ from the other species by the greater variability in the number of vein pairs per leaf, with a range of 3 – 9 pairs (vs 3 – 5 pairs). The abaxial leaf surface and fruit of *S. velutina*, however, are significantly more densely hairy than in *S. macrocarpa*. Otherwise, both species are similar morphologically, as noted by Cahen & Utteridge (2017a).

Clade C (*S. lanceata*) is sister to the clade containing Clades A and B. This species has unique fruit and leaf characteristics among the sampled Ventilagineae. The apical fruit appendage is enlarged and inflated. This species is mostly coastal, and fruit have been observed floating in seawater (Ridley 1930). Instead of domatia as tufts of trichomes, a character uniting the sister group, *S. lanceata* has pockets formed by extensions from the leaf veins. Like the domatia of other species, these are positioned at the intersection between the midrib and the primary veins.

Clade D (*V. denticulata* and *V. madraspatana*) is sister to the clade containing Clades A, B, and C. They are united by the characters of having seed chambers meeting the wing at a sharp angle and the absence of domatia. *Ventilago denticulata* differs from *V. madraspatana* in the calyx covering c. 50% of the seed chamber (vs under 30% in *V. madraspatana*, often much less). In addition, the leaves of *V. denticulata* tend to be larger, c. 10 × 5 cm (vs c. 6 × 3 cm), the flower disks densely hairy (vs sparsely hairy), and the indumentum of the young leaves, branchlets and inflorescence bright yellow (vs often a more brownish yellow) (D. Cahen, pers. obs.). Most characters that are shared between *V. denticulata* and *V. madraspatana* and their sister clades (Clade A, B and C) are also present in all the sampled Ventilagineae species.

Clade E (*V. viminalis* and *V. leiocarpa*) is sister to all the previously mentioned species. They share the characters of having similar fruit dimensions (length/width), a small proportion of fruit chamber enclosed in the calyx, and glabrous fruit. A clear difference is that *V. viminalis* after developing as a scrambling plant, becomes a fully self-supporting tree by maturity, the only known case in Ventilagineae (Kellermann & Thiele 2024). The species also has unique linear leaves, more than 5 times longer than wide. *Ventilago leiocarpa* is one of a few species in the genus—excluding *Smythea*—that have persistent leaves subtending fascicles of open flowers and fruits. Other species with this character include *V. buxoides* Baill. and *V. elegans* Hemsl., which were not included in this analysis. No morphological characters have yet been identified that unite *V. viminalis* and *V. leiocarpa* and distinguish them from their sister species.

Clade F (*V. malaccensis*, *V. dichotoma*, and *V. maingayi*) is sister to the clade containing Clades A–E. These species are united in having glabrous fruit. Within this clade, *V. malaccensis* and *V. dichotoma* are sister to *V. maingayi*, which differs in having an unusually high number of leaf vein pairs to a maximum of c. 18. This contrasts with 3–9 in the sister group. *Ventilago malaccensis* has a cup-like calyx that covers c. half of the seed chamber, unlike *V. dichotoma*, which has an annular calyx attached at the base of the seed chamber. For additional characters separating these two species, see the Key to *Ventilago* species of Borneo (Cahen & Utteridge 2017b).

## Discussion

We sequenced several species of *Smythea* for the first time to answer key taxonomic questions. Our results support the decision to sink *Smythea* into *Ventilago*, which was initially indicated by the recent phylogeny of Tian *et al*. (2024), showing that *Ventilago* is paraphyletic with *Smythea* nested within it. Given the paraphyly of *Ventilago*, we considered two options. One possibility was preserving the genus *Smythea* through further splitting of *Ventilago.* This would create several smaller genera whilst preserving monophyly. Another option was to transfer all *Smythea* species to the genus *Ventilago* by making new combinations.

The option of describing segregate genera would have the advantage of maintaining the genus *Smythea*, distinguished morphologically from *Ventilago* by the tapering seed chamber. This option would avoid the need for revision of previously published accounts including the Flora of Thailand (Norsaengsri *et al*. 2020), which treat *Smythea* and *Ventilago* as distinct genera. However, retaining the name *Smythea* would necessitate describing segregate genera of *Ventilago* due to its paraphyly. This would be challenging due to the lack of discrete morphological features associated with the *Ventilago* clades (the candidates for new genera). This would present difficulties in identifying the correct generic placement for the many *Ventilago* species that have not been included in the current phylogeny.

The alternative option is to merge *Smythea* with *Ventilago*, formally synonymising the genera. Synonymising *Smythea* with *Ventilago* involves making new combinations, and restoring the names of other *Smythea* species originally published in *Ventilago*. The authors conclude that until a phylogenetic analysis including a higher proportion of *Ventilago* species is carried out, making taxonomic changes beyond sinking *Smythea* into *Ventilago* would not be justified.

Despite the disruption that could be caused by synonymising the genera, the authors believe this would be considerably less than the alternative option discussed of describing segregate genera. For this reason, we have proceeded with synonymising *Smythea* with *Ventilago*.

The expanded concept of *Ventilago* is equivalent to the tribe Ventilagineae. It is morphologically distinguishable from all other genera in the Rhamnoid group by the presence of a pronounced apical appendage in the form of a wing or air-filled pocket and is readily identified from herbarium specimens as well as in the field. The addition of species previously described as *Smythea* brings the total number of species from 40 to 52 (Plants of the World Online 2025).

These phylogenetic results we have obtained are broadly consistent with morphology. For example, *Smythea lanceata* (= *V. lanceata*), the only species sampled with inflated seed chambers, is sister to the rest of *Smythea*. It is possible that *V. poomae*, a species known from one collection and also with an inflated seed chamber, would form a clade with *V. lanceata*. However, *V. poomae* is an inland species whilst *V. lanceata* is mostly coastal and the two species have non-overlapping ranges. It is also possible that the inflated fruit evolved independently in these species.

The phylogenetic tree generated for this study sheds light on aspects of the evolution of *Ventilago*. For instance, the water-dispersed *V. lanceata* is nested within a clade of wind dispersed *Ventilago*, indicating that their common ancestor was wind dispersed.

The Australian endemic *V. viminalis*, is nested within a clade containing Asian species all with a climbing habit. This demonstrates that the tree habit evolved from climbers and indicates that the genus diversified in Asia before reaching Australia.

The oldest known fossil of *Ventilago,* and the only dating from the Eocene, is a specimen of *Ventilago tibetensis* C.-Del Rio & T.-Su, described based on fossil fruit. The fossil was collected from central Tibet and estimated to be at least 47 million years old (Del Rio *et al*. 2021). There is also an Oligocene fossil from Mexico of the extinct species *Ventilago engoto* Calvillo-Canadell and Cevallos-Ferriz (Calvillo Canadell *et al*. 2007). Several more recent fossils from the Miocene have been found in central Asia. These include specimens of the extinct species *V. lincangensis* K. N. Liu & S. P. Xie, *V. ovatus* Konomatsu & Awasthi and *V. tistaensis* Antal & Prasad, as well as those resembling the extant species *V. madraspatana*. Based on evaluation of the fossil record, Del Rio *et al*. suggest that the Indo-Tibetan region may be the origin of the genus, and that the location of the Oligocene fossil from Mexico may be explained by dispersal via the Bering Land Bridge during the Eocene (Del Rio *et al*. 2021).

In an earlier publication, Richardson *et al*. (2000a) propose an alternative scenario, in which *Ventilago* originated in Gondwana and then spread into Asia when India collided with Asia, with India serving as a raft. This theory known as the “Out-of-India” hypothesis has been investigated in other Rhamnaceous genera such as *Paliurus* (Chen *et al*. 2017). Our analysis did not include any African species of *Ventilago*, so we cannot conclusively test this hypothesis. However, we note that the Indian species of *Ventilago*, *V. bombaiensis*, *V. denticulata*, and *V. madraspatana*, are all deeply nested within the genus, whereas Malesian species, such as *V. dichotoma*, *V. maingayi*, and *V. malaccensis*, are sister to the rest of the genus. This suggests that the genus may have diversified, and possibly originated, in Malesia before dispersing and diversifying in other regions. This pattern is similar to that of *Sageretia* Brongn. in the Rhamnaceae, which was reconstructed by Yang *et al*. (2019) as having an origin in tropical Asia, followed by a migration to more temperate parts of Asia, Africa via Arabia, and America via the Bering Land Bridge. More sampling is necessary to better understand the evolutionary history of the genus. We included only 13 of over 50 known species of *Ventilago* (including *Smythea*) in the molecular phylogeny. A more complete sampling of the genus, using a molecular clock calibrated with the known fossils and combined with a biogeographical analysis, would improve inferences on the present-day distribution of the genus. This approach has been recently applied to other genera of the Rhamneae tribe, such as *Berchemia* (Huang *et al*. 2021) and *Sageretia* (Yang *et al*. 2019).

A phylogenetic analysis is only as robust as the level of resolution of the underlying taxonomy, and much work remains to be done to characterise and describe species before *Ventilago* can be considered taxonomically well understood. Except for the *Flora of Thailand* (Norsaengsri *et al*. 2020), no comprehensive treatment of *Ventilago* is available in the major flora treatments of Southeast Asia. Species concepts of *Ventilago* in *Flora of Thailand* may need to be reevaluated after further analysis, particularly for the set of specimens with glabrous fruit wings and calyx remains that cover less than a quarter of the seed chamber at the base of mature fruits (Cahen 2022). Recent contributions helped to better understand the diversity of Ventilagineae (Cahen & Utteridge 2017, 2018; Cahen *et al*. 2020; Utteridge & Cahen 2021). However, work is still required to distinguish groups of morphologically similar species. This includes a group of species of *Ventilago* with broad leaves with entire margins, 6–8 pairs of abaxially almost flat secondary veins, no domatia at secondary vein axils, and flowers with hairy nectary discs (*V. papuana* Merr. & L.M.Perry, *V. borneensis* Ridl., *V. microcarpa* K.Schum. and *V. madraspatana* sensu Gaertn.) (Cahen 2022). There is also a set of *Ventilago* specimens with glabrous fruit wings and calyx remains covering less than a quarter of the seed chamber at the base of mature fruits that seem to form a morphological continuum without well-defined and clearly identifiable species (*V. maingayi*, *V. dichotoma*, *V. harmandiana* Pierre, *V. sororia* Hance, etc.) (Cahen 2022); whether these should be recognised as separate or conspecific should be investigated.

Producing a well-supported phylogenetic tree of *Ventilago* aids *in-situ* conservation as a tree would provide evidence to sink, erect or maintain taxa and thus clarify the identity of the entities that should be the focus of conservation efforts. Both *V. calpicarpa* Kurz and *V. poilanei* are known from only one and two collections, respectively (Cahen & Utteridge, 2017b). This lack of data results in lower certainty of the legitimacy of these taxa, as *Ventilago* species are known to have variable morphology (H. Miller, pers. obs.). However, if molecular data supports these recently described species, this increases their legitimacy. With more stable species concepts, IUCN Red List assessments published will be less likely to need to be revised as specimens are redetermined. So far, only preliminary conservation assessments have been made based on species described using morphology within *Ventilago* (Cahen & Utteridge 2017a,b; Cahen *et al*. 2020; Utteridge & Cahen 2021). If the Red List statuses of *Ventilago* species are known, this can inform the establishment of protected areas through schemes such as the New Guinea Tropical Important Plant Areas (TIPAs) project, which identifies areas with exceptional diversity and the presence of large numbers of threatened taxa. This allows prioritisation of sites for conservation and management through collaboration with partner countries (Royal Botanic Gardens, Kew 2022).

### New combinations in *Ventilago*

Based on the evidence of the molecular tree presented, we propose to merge *Smythea* into *Ventilago* formally synonymising the genera and producing seven new combinations. In addition, the five remaining *Smythea* species, which have previously been placed in *Ventilago* are restored to *Ventilago*. Full synonymy for *Smythea* species is available in Cahen & Utteridge (2017a). The new and restored combinations are as follows:

1. ***Ventilago batanensis*** (Cahen & Utteridge) Cahen, Utteridge & Henry Mill. **comb. nov.** [IPNI Code] ≡ *Smythea batanensis* Cahen & Utteridge, *Kew Bull.* 73(1)-2: 6 (2017a). Type: Philippines, Batan Islands, Batanes, Mt Iraya [20°28’N 122°00’E], May 1930, Ramos BS 80170 (holotype K [K000606763]; isotype K [K000606764]). **DISTRIBUTION.** Endemic to the Philippines.
2. ***Ventilago beccarii*** (Cahen & Utteridge) Cahen, Utteridge & Henry Mill. **comb. nov.** [IPNI Code] ≡ *Smythea beccarii* Cahen & Utteridge, Kew Bull. 73(1)-2: 8 (2017a). Type: Indonesia, Sulawesi, Kandari [Kendari] [4°3’S 122°32’E], 1874, Beccari s.n. (holotype K [K000606778], isotypes FI, K [K000606779]). **DISTRIBUTION.** Endemic to Sulawesi.
3. ***Ventilago hirtella*** (Cahen & Utteridge) Cahen, Utteridge & Henry Mill. **comb. nov.** [IPNI Code] ≡ *Smythea hirtella* Cahen & Utteridge, Kew Bull. 73(1)-2: 12 (2017a). Type: Malaysia, Sabah, Lahad Datu, Tabin Wildlife Reserve, 19 June 2000, Madani *et al*. 145373 (holotype K! [K000271426]; isotype SAN). **DISTRIBUTION.** Endemic to Borneo.
4. ***Ventilago macrocarpa*** (Hemsl.) Cahen, Utteridge & Henry Mill. **comb. nov.** [IPNI Code] ≡ *Smythea macrocarpa* Hemsl., Hooker’s Icon. Pl. 16: t. 1558 (1886). Type: Malaysia, Peninsular Malaysia, Perak, Larut [Taiping], Waterfall Hill, s.a., Wray 36 (lectotype K [K000681974]; isolectotype: K [K000681973]). **DISTRIBUTION.** Asia: Borneo, Malaya, Sumatra, Thailand.
5. ***Ventilago papuana*** (Utteridge & Cahen) Cahen, Utteridge & Henry Mill **comb. nov.** [IPNI Code] ≡ *Smythea papuana* Utteridge & Cahen, Phytotaxa 498(3): 153 (2021). Type: Papua New Guinea. Northern Div. [Oro Province], ca 1 km N of Oitatandi village [8°36’S 147°57’E], alt. ca 25 m, 19 Aug. 1953 (fr.), Hoogland 3685 (holotype: CANB acc. no. 74365; isotype: LAE n.v.). **DISTRIBUTION.** Endemic to New Guinea.
6. ***Ventilago poilanei*** (Cahen & Utteridge) Cahen, Utteridge & Henry Mill. comb. **nov.** [IPNI Code] ≡ *Smythea poilanei* Cahen & Utteridge, Kew Bull. 73(1)-2: 21 (2017a). Type: Laos, Vientiane Prefecture, Ban Tha Ngon Road [18°5’N 102°40’E], 170 m, 1 Oct. 1955, de Malahide 88 (holotype K [K000606765]; isotypes K [K000606766], SING). **DISTRIBUTION.** Endemic to Laos.
7. ***Ventilago poomae*** (Cahen & Utteridge) Cahen, Utteridge & Henry Mill. **comb. nov.** [IPNI Code] ≡ *Smythea poomae* Cahen & Utteridge, Kew Bull. 73(1)-2: 21 (2017a). Type: Thailand, Nan Province, Pua, Doi Phu Kha National Park [19°11’55’’N 101°4’ 46’’E], 1700 m, 17 Aug. 1995, Pooma 1112 (holotype BKF! [BKF 102045]). **DISTRIBUTION.** Endemic to Thailand. <u>Resurrected names in *Ventilago*</u> The following species, which were originally published in *Ventilago* but subsequently transferred to *Smythea*, are returned to *Ventilago*:
8. ***Ventilago bombaiensis*** Dalzell, Hooker’s J. Bot. Kew Gard. Misc. 3: 36 (1851). ≡ *Smythea bombaiensis* (Dalzell) Banerjee & P.K.Mukh., Indian Forester 96: 214 (1970).
9. ***Ventilago calpicarpa*** (Kurz) Oza, Indian Forester 94: 267 (1968) ≡*Smythea calpicarpa* Kurz, J. Asiat. Soc. Bengal, Pt. 2, Nat. Hist. 41: 301 (1872)
10. ***Ventilago lanceata*** Tul, Ann. Sci. Nat., Bot., sér. 4, 8: 121 (1857) _≡_*Smythea lanceata* (Tul.) Summerh., in Bull. Misc. Inform. Kew 1928: 389 (1928)
11. ***Ventilago oblongifolia*** Blume, Bijdr. Fl. Ned. Ind.: 1144 (1826) _≡_*Smythea oblongifolia* (Blume) Cahen & Utteridge, Kew Bull. 73(1)-2: 18 (2017)
12. ***Ventilago velutina*** Ridl, Fl. Malay Penins. 1: 467 (1922) _≡_*Smythea velutina* (Ridl.) S.P.Banerjee & P.K.Mukh. Bull. Bot. Surv. India 10: 251 (1969) <u>Typification of *Ventilago* and synonymisation of *Smythea* with *Ventilago*:</u> **Ventilago** Gaertn. (Gaertner 1788: 223). Type species: *Ventilago madraspatana* Gaertn. (Gaertner 1788: 223). Type: India, Mysore, Hassan District, Shiradi ghat, 12 April 1969, *Saldanha* 13285 (neotype, selected here: K! [K004289083]). *Smythea* Seem. (Seeman 1862: 69), **synon. nov.** Type species: *Smythea pacifica* Seem. (Seeman 1861: 255). Type: Fiji, Viti Levu, June 1860, *Seemann* 79 (lectotype, designated by Cahen & Utteridge (2018): K! [K000681971]; isolectotypes BM!-image seen [BM000838644], P!-image seen [P06886654]) = *Ventilago lanceata* Tul.

When reviewing the taxonomy of the names for this study it was clear a type specimen was required for *Ventilago*. In his protologue, Gaertner (1788) referenced *Funis viminalis* from Ambon Island in the Maluku Islands of Indonesia, described by Rumphius (1747) (“*Funis viminalis*. RUMPH. *amb*. 5.p. 3 t. 2.”), and unspecified material from Joseph Banks’s herbarium (“E collect. Banksiana”).

As discussed by Cahen & Utteridge (2018) and Cahen (2022), issues arise regarding the application of the name *Ventilago madraspatana*, as material from Ambon belongs to a distinct species separate from the Indian material typically identified as *V. madraspatana*. Cahen (2022) noted that this issue had not been formally resolved. However, instead of Gaertner’s name being misapplied, it is more likely that he examined Indian material alongside the Ambonese species—potentially the unspecified specimens from Banks’s herbarium. His use of the epithet *madraspatana* strongly suggests connections to collections near Chennai (formerly Madras).

It appears in this case that Gaertner applied a broad species concept, encompassing both Ambonese and Indian material. The illustration in Rumphius’s *Herbarium amboinense* could not be selected as the lectotype because it presumably depicts the Ambonese species. Additionally, the material from Banks’s collection could not be located. To avoid confusion and considering that the name *V. madraspatana* has been widely applied to the Indian species, a neotype is designated here. Therefore, *Saldanha* 13285 from Hassan District, Karnataka, is selected as the neotype because it clearly represents the species in its widely accepted concept, with both leaf surfaces visible, several mature fruits, and has precise collection data. The neotype was collected just 450 km from Chennai, whereas Ambon is located over 5,000 km away.

## Acknowledgements

We would like to acknowledge Jürgen Kellermann and one anonymous reviewer for their comments which have greatly improved our article. The first author would also wish to thank László Csiba for his assistance with training in DNA extractions at the Jodrell Laboratory.

## Ethics declarations

### Data availability

Raw sequence data are deposited in Sequence Read Archive (SRA) available on NCBI servers under BioProject number PRJNA1100519 (http://www.ncbi.nlm.nih.gov/bioproject/PRJNA1100519). All scripts related to phylogenetic analyses follow Sumanon *et al*. 2023 and are available on GitHub at https://github.com/pebgroup/Maesa_sptree

The alignments, gene trees, and species trees are deposited on Zenodo at 10.5281/zenodo.10976870.

### Author contributions

DC provided supervision during the development of this paper as an MSc project and reviewed the manuscript especially the introductory and taxonomy section. TU also conceived of and developed this project and fully reviewed the manuscript. NM completed laboratory work for illumina sequencing at Kew’s Jodrell Laboratory. PS completed phylogenetic analysis and produced phylogenetic trees. FF provided comments on the manuscript, especially the phylogenetics section. Sampling and DNA extractions were completed by HM as well as the writing of the manuscript.

---

**Appendix 1.** list of species, specimens and SRA accession numbers of data used in this study. All read sequences, except of the outgroup (*Maesopsis eminii*), were newly generated and deposited on Sequence Read Archive (SRA: https://www.ncbi.nlm.nih.gov/sra) data of NCBI server.

| Taxon name | Tip label in the phylogeny | Collector and collector number (herbarium code) | Locality | Date of collection |
| --- | --- | --- | --- | --- |
| *Ventilago bombaiensis* | *Smythea bombaiensis* | Stocks s.n. (K) | India | s.a. |
| *Ventilago lanceata* | *Smythea lanceata* | 10807 (K) | Malaysia, Jahore | 13/6/1965 |
| *Ventilago macrocarpa* | *Smythea macrocarpa* | De Wilde 13903 (K) | North Sumatra | 30/7/1972 |
| *Ventilago oblongifolia* | *Smythea oblongifolia* 1 | 168 (K) | Indonesia, Timor Island | 11/8/2011 |
| *Ventilago oblongifolia* | *Smythea oblongifolia* 2 | A.Z 3840 (K) | Malaysia, Perlis state | 23/8/1991 |
| *Ventilago poilanei* | *Smythea poilanei* | Malahyde 88 (K) | Laos | 1/10/1955 |
| *Ventilago velutina* | *Smythea velutina* 1 | 1984 (K) | Malaysia, Jelubu District | 9/6/1996 |
| *Ventilago velutina* | *Smythea velutina* 2 | T & P 402 (K) | Malaysia, Ulu Klang Forest reserve | 10/7/1971 |
| *Ventilago denticulata* | *Ventilago denticulata* | Larsen 44384 (K) | Thailand | 15/11/1993 |
| *Ventilago dichotoma* | *Ventilago dichotoma* | Andau 602 (K) | Malaysia | 3/9/1996 |
| *Ventilago leiocarpa* | *Ventilago leiocarpa* | Hu, S.Y. 11890 (K) | Hong Kong, China | 19/4/1972 |
| *Ventilago madraspatana* | *Ventilago madraspatana* | Saldanha 13285 (K) | India, Hassan District | 12/4/1969 |
| *Ventilago maingayi* | *Ventilago maingayi* | Matasri 26867 (K) | Malaysia, Kepong Selangor | 7/7/1993 |
| *Ventilago malaccensis* | *Ventilago malaccensis* | T & P 257 (K) | Malaysia | 7/5/1969 |
| *Ventilago viminalis* | *Ventilago viminalis* 1 | Kenneally, K.F. 9507 (K) | Western Australia | 28/8/1985 |
| *Ventilago viminalis* | *Ventilago viminalis* 2 | GAQLD0252 (K) | Queensland Australia | 30/11/2006 |

